## Supplementary Figures 1-13 for "Humidity-driven ABA depletion determines plant-pathogen competition for leaf water"

**Supplementary materials**

Supplementary Figures 1-13

Supplementary Tables 1-7

Supplementary Movie 1

**Supplementary Fig. 1: Characterization of *cyp707a* mutant plants.**

**a**, Five-week-old plants grown under short-day conditions. Scale bars = 3 cm. **b**, Genotyping of the indicated plants by PCR. LB-RP and LP-RP primer pairs specifically amplify mutated and non-mutated DNA, respectively.

**Supplementary Fig. 2: Characterization of *cyp707a1 cyp707a3 ost2-3D* mutant plants.**

**a**, Five-week-old plants grown under short-day conditions. Scale bars = 3 cm. **b**, Genotyping of the indicated plants by PCR and Sanger sequencing. LB-RP and LP-RP primer pairs specifically amplify mutated and non-mutated DNA, respectively.

**Supplementary Fig. 3: ABA depletion enhances bacterial resistance by restricting water-soaking independently of SA under high humidity.**

**a**, **d**, Five-week-old plants grown under short-day conditions. Scale bars = 3 cm. **b**, **e**, Genotyping of the indicated plants by PCR. LB-RP and LP-RP primer pairs specifically amplify mutated and non-mutated DNA, respectively. **c**, **f,** Water-soaking in *Pst* Δ*hrcC*- (**c**) or *Pst* DC3000-inoculated (**f**) leaves. The indicated plants were infiltrated with *Pst* Δ*hrcC* or *Pst* DC3000 ($OD_{600}$ = 0.02) and maintained under high humidity (HH) for 1 day. The number of water-soaked leaves was pooled from two independent experiments. **g**, **h**, Bacterial growth in *Pst* DC3000-inoculated leaves. The indicated plants were infiltrated with *Pst* DC3000 ($OD_{600}$ = 0.0002) and maintained under HH (**g**) or moderate humidity (MH) (**h**) for 3 days. Bars represent means ($n$ = 6). Different letters indicate statistically significant differences ($P < 0.05$; one-way ANOVA followed by Tukey's test). WT, wild-type; dpi, days post-infiltration. Experiments in (**g**) and (**h**) were repeated twice with similar results.

**Supplementary Fig. 4: CYP707A3/A1-mediated ABA depletion and PTI contribute**

**additively to bacterial resistance under high humidity.**

**a**, Expression levels of *FRK1*, *NHL10*, and *PHI-1* in *Pst* Δ*hrcC*-infiltrated leaves under different humidity conditions. The indicated plants were maintained under moderate humidity (MH) or high humidity (HH) for 24 h before infiltration with water (Mock) or *Pst* Δ*hrcC* ($OD_{600}$ = 0.2). Infiltrated plants were kept under the same humidity conditions for 1 h. Bars represent means ($n$ = 3). **b**, Five-week-old plants grown under short-day conditions. Scale bars = 3 cm. **c**, Genotyping of the indicated plants by PCR and Sanger sequencing. LB-RP and LP-RP primer pairs specifically amplify mutated and non-mutated DNA, respectively. **d,** Water-soaking in *Pst* Δ*hrcC*-infiltrated leaves. The indicated plants were infiltrated with *Pst* Δ*hrcC* ($OD_{600}$ = 0.02) and maintained under HH for 1 day. The number of water-soaked leaves was pooled from two independent experiments. **e**, Bacterial growth in *Pst* Δ*hrcC*-infiltrated leaves. The indicated plants were infiltrated with *Pst* Δ*hrcC* ($OD_{600}$ = 0.02) and maintained under HH for 3 days. Bars represent means ($n$ = 6). Different letters indicate statistically significant differences ($P <$ 0.05; two-way ANOVA followed by Tukey's test in **a**; one-way ANOVA followed by Tukey's test in **e**). WT, wild-type; *bbc*, *bak1-5 bkk1 cerk1*; dpi, days post-infiltration. Experiments in (**a**) and (**e**) were repeated twice with similar results.

**Supplementary Fig. 5: Gene Ontology analysis of high humidity-induced genes.**

Top 10 enriched Gene Ontology categories of humidity-induced genes in the clusters identified in Fig. 3c.

**Supplementary Fig. 6: The CGCG boxes in the *CYP707A3* promoter are critical for water-soaking resistance.**

**a**, Sequence of the *CYP707A*3 promoter region showing the deleted CGCG boxes in the ΔCGCG construct. Numbers indicate the position relative to the ATG start codon. **b**, Five-week-old plants grown under short-day conditions. Scale bars = 3 cm. **c**, Genotyping of the indicated plants by PCR. LB-RP and LP-RP primer pairs specifically amplify mutated and non-mutated DNA, respectively. **d**, Expression levels of *CYP707A3-FLAG* in humidity-treated leaves. The indicated plants were exposure to moderate humidity (MH) or high humidity (HH) for 0.5 h. Bars represent means ($n$ = 3). Different letters indicate statistically significant differences ($P < 0.05$; two-way ANOVA followed by Tukey's test).

**e**, Water-soaking in *Pst* Δ*hrcC*-inoculated leaves. The indicated plants were infiltrated with *Pst* Δ*hrcC* ($OD_{600}$ = 0.02) and maintained under HH for 1 day. The number of water-soaked leaves was pooled from three independent experiments. WT, wild-type; dpi, days post-infiltration.

**Supplementary Fig. 7: High humidity-triggered cytosolic $Ca^{2+}$ elevation precedes *CYP707A3* induction.**

**a**, **b,** Cytosolic $Ca^{2+}$ levels in humidity-treated leaves. Two-week-old *35S::GCaMP3* transgenic plants grown under long-day conditions were exposed to moderate humidity (MH) or high humidity (HH) for the indicated time, and GCaMP3 fluorescence signals were measured. Representative GCaMP3 fluorescence images from HH-treated plants are shown in (**a**), and quantified GCaMP3 fluorescence signals are presented in (**b**). Bars represent means ± SEM ($n$ = 10). **c**, Effect of $Ca^{2+}$ channel inhibition on high humidity-induced *CYP707A3* and *CYP707A1* expression. Wild-type (WT) plants were infiltrated with the indicated inhibitors, maintained under MH for 24 h, and then exposed to MH or HH for 0.5 h. Bars represent means ($n$ = 3). **d**, Temporal dynamics of *CYP707A3* expression in high humidity-treated shoots. Two-week-old WT plants grown under long-day conditions were exposed to HH for the indicated time. Bars represent means ($n$ = 3). Different letters indicate statistically significant differences ($P < 0.05$; two-way ANOVA followed by Tukey's test in **c**; one-way ANOVA followed by Tukey’s test in **d**). Experiments in (**c**) and (**d**) were repeated twice with similar results.

**Supplementary Fig. 8: High humidity-induced *CYP707A3* expression is impaired in *cngc2*, *cngc4*, and *cngc9* mutant plants.**

**a**, **b**, Expression levels of *CYP707A3* in humidity-treated shoots. The indicated plants were grown under long-day conditions for 2 weeks and then exposed to moderate humidity (MH) or high humidity (HH) for 0.5 h. Results from the first experiments are shown in (**a**), and results from the second experiments are shown in (**b**). Bars represent means ($n$ = 2-3). Different letters indicate statistically significant differences ($P < 0.05$; two-way ANOVA followed by Tukey's test). WT, wild-type.

**Supplementary Fig. 9: Characterization of *camta3* mutant plants.**

**a**, Five-week-old plants grown under short-day conditions. Scale bars = 3 cm. **b**, Genotyping of the indicated plants by PCR and Sanger sequencing. LB-RP and LP-RP primer pairs specifically amplify mutated and non-mutated DNA, respectively. **c**, SA and JA-Ile levels in humidity-treated leaves. The indicated plants were exposed to moderate humidity (MH) or high humidity (HH) for 2 h. Bars represent means ($n$ = 5). Different letters indicate statistically significant differences ($P$ < 0.05; two-way ANOVA followed by Tukey's test). WT, wild-type; CAMTA3-A855V, a partial loss-of-function variant.

**Supplementary Fig. 10: CAMTA3 protein levels in *Pst* DC3000-inoculated leaves.**
Wild-type (WT) and transgenic plants expressing CAMTA3-GFP driven by the *CAMTA3* promoter (*pCAMTA3::CAMTA3* in *camta2/3* #3) were infiltrated with the indicated *Pst* DC3000 strains ($OD_{600}$ = 0.02) and maintained under moderate humidity (MH) for 8 h. CAMTA3-GFP was detected by immunoblotting with anti-GFP antibody (α-FLAG). The ponceau-stained blot is shown as a loading control. The experiment was repeated three times with similar results.

**Supplementary Fig. 11: Characterization of *DEX::AvrPtoB*, *DEX::AvrPto*, *and* *DEX::AvrPtoB* transgenic plants.**
**a**, Five-week-old plants grown under short-day conditions. Scale bars = 3 cm. **b**, Expression levels of AvrPtoB-FLAG and AvrPto-FLAG in DEX-treated leaves. The indicated plants were infiltrated with 0.1% ethanol (−DEX ) or 10 µM DEX (+DEX) and maintained under moderate humidity (MH) for 24 h. AvrPtoB-FLAG and AvrPto-FLAG were detected by immunoblotting with anti-FLAG antibody (α-FLAG). The ponceau-stained blot is shown as a loading control. **c-e**, Expression levels of *CYP707A1* in humidity-treated leaves with or without *AvrPtoB* (**c**), *AvrPto* (**d**), and *AvrPtoB-F479A* (**e**). The indicated plants were treated with DEX as described in (**b**), followed by exposure to MH or high humidity (HH) for 0.5 h. Bars represent means ($n$ = 3). Different letters indicate statistically significant differences ($P$ < 0.05; two-way ANOVA followed by Tukey's test). The experiments in (**c**) and (**d**) were repeated twice with similar results.

**Supplementary Fig. 12: *Pst* DC3000-mediated *CYP707A3* suppression and ABA accumulation are compromised by loss of *avrPtoB* and *avrPto***

**a**, Schematic overview of the experimental design for monitoring temporal dynamics of *CYP707A3* expression and ABA accumulation in *Pst* DC3000-inoculated leaves under high humidity (HH). **b**, Temporal dynamics of *CYP707A3* expression in *Pst* DC3000-inoculated leaves. Wild-type (WT) plants were infiltrated with the indicated *Pst* DC3000 strains ($OD_{600}$ = 0.2), maintained under moderate humidity (MH) for 2 h (2 hours post-infiltration, hpi), and then transferred to HH for 4 h (6 hpi), 10 h (12 hpi), or 22 h (24 hpi). Bars represent means ($n$ = 3). **c**, Temporal dynamics of ABA levels in *Pst* DC3000-inoculated leaves. WT plants were infiltrated with the indicated *Pst* DC3000 strains as described in (**a**) and (**b**). Bars represent means ($n$ = 4). Different letters indicate statistically significant differences ($P < 0.05$; two-way ANOVA followed by Tukey's test). Experiment in (**b**) was repeated twice with similar results.

**Supplementary Fig. 13: Characterization of *Pst* Δ*avrPtoB* and *Pst* Δ*avrPto* mutant strains.**

**a, b**, Generation of *Pst* DC3000 mutant strains lacking *AvrPtoB* (Δ*avrPtoB*) or *AvrPto* (Δ*avrPto*). Deleted genomic regions (**a**) and genotyping results (**b**) of the established *Pst* DC3000 mutants. Numbers indicate the position relative to the ATG start codon. Arrows indicate genotyping primers. M, DNA size marker. **c**, **d**, Representative images (**c**) and quantification (**d**) of water-soaking in *Pst* DC3000-inoculated leaves. Wild-type (WT) plants were infiltrated with the indicated *Pst* DC3000 strains ($OD_{600}$ = 0.2), maintained under MH for 2 h, and then transferred to HH for 20-22 h. Bars represent the median ($n$ = 36; pooled from two independent experiments). Different letters indicate statistically significant differences ($P < 0.05$; one-way ANOVA followed by Tukey's test).

**Supplementary Table 1**

Detailed information of DEGs in Fig. 3a.

**Supplementary Table 2**

Detailed information of 1000 most variable genes in Fig. 3c.

**Supplementary Table 3**

Detailed information of 2000 most variable genes in Fig. 5e.

**Supplementary Table 4**

List of Arabidopsis plants used in this study.

**Supplementary Table 5**

List of plasmid DNA used in this study.

**Supplementary Table 6:**

List of primers used in this study.

**Supplementary Table 7**

List of accession numbers related to this study.

**Supplementary Movie 1:**

Cytosolic $Ca^{2+}$ dynamics in leaves upon exposure to high humidity. Arabidopsis transgenic plants expressing *35S::GCaMP3* were exposed to high humidity, and cytosolic $Ca^{2+}$ changes were visualized through GCaMP3 fluorescence. Signals were captured at 0.08 min intervals over the indicated time course.

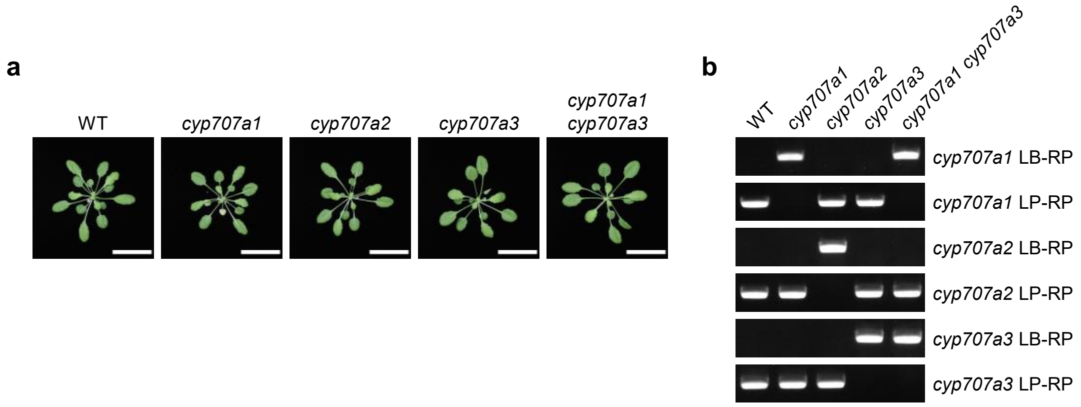


**Supplementary Fig. 1: Characterization of *cyp707a* mutant plants.**
**a**, Five-week-old plants grown under short-day conditions. Scale bars = 3 cm. **b**, Genotyping of the indicated plants by PCR. LB-RP and LP-RP primer pairs specifically amplify mutated and non-mutated DNA, respectively.

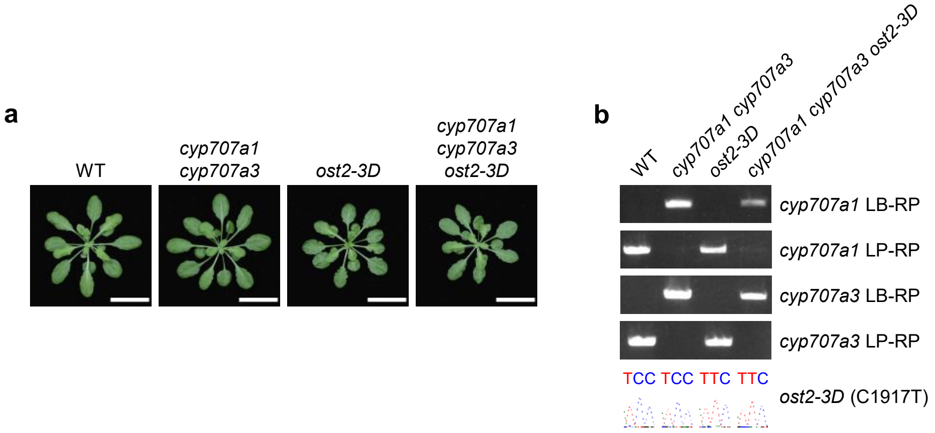


**Supplementary Fig. 2: Characterization of *cyp707a1 cyp707a3 ost2-3D* mutant plants.**
**a**, Five-week-old plants grown under short-day conditions. Scale bars = 3 cm. **b**, Genotyping of the indicated plants by PCR and Sanger sequencing. LB-RP and LP-RP primer pairs specifically amplify mutated and non-mutated DNA, respectively.

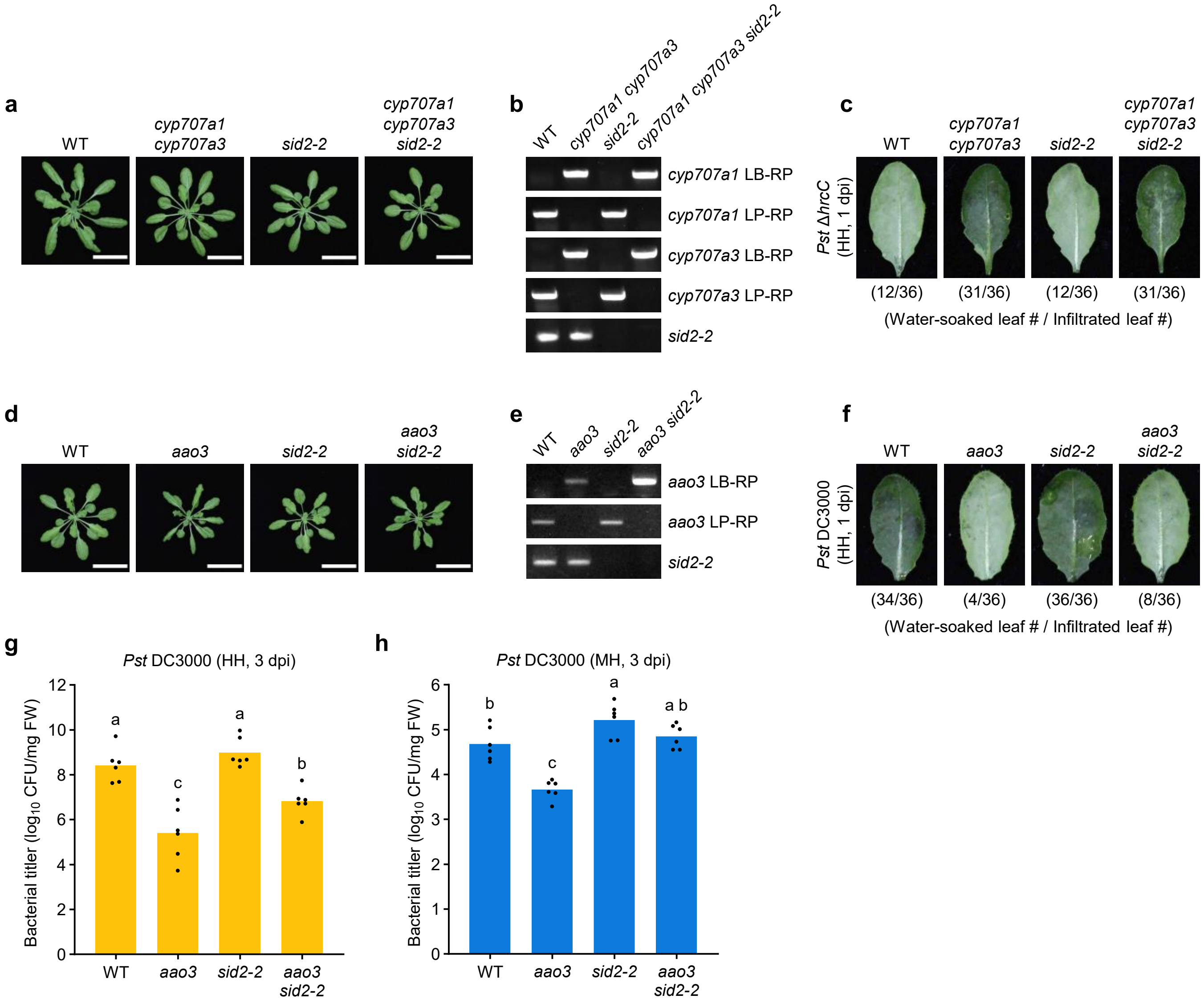


**Supplementary Fig. 3: ABA depletion enhances bacterial resistance by restricting water-soaking independently of SA under high humidity.**

**a**, **d**, Five-week-old plants grown under short-day conditions. Scale bars = 3 cm. **b**, **e**, Genotyping of the indicated plants by PCR. LB-RP and LP-RP primer pairs specifically amplify mutated and non-mutated DNA, respectively. **c**, **f**, Water-soaking in *Pst* Δ*hrcC*- (**c**) or *Pst* DC3000-inoculated (**f**) leaves. The indicated plants were infiltrated with *Pst* Δ*hrcC* or *Pst* DC3000 ($OD_{600}$ = 0.02) and maintained under high humidity (HH) for 1 day. The number of water-soaked leaves was pooled from two independent experiments. **g**, **h**, Bacterial growth in *Pst* DC3000-inoculated leaves. The indicated plants were infiltrated with *Pst* DC3000 ($OD_{600}$ = 0.0002) and maintained under HH (**g**) or moderate humidity (MH) (**h**) for 3 days. Bars represent means ($n$ = 6). Different letters indicate statistically significant differences ($P < 0.05$; one-way ANOVA followed by Tukey's test). WT, wild-type; dpi, days post-infiltration. Experiments in (**g**) and (**h**) were repeated twice with similar results.

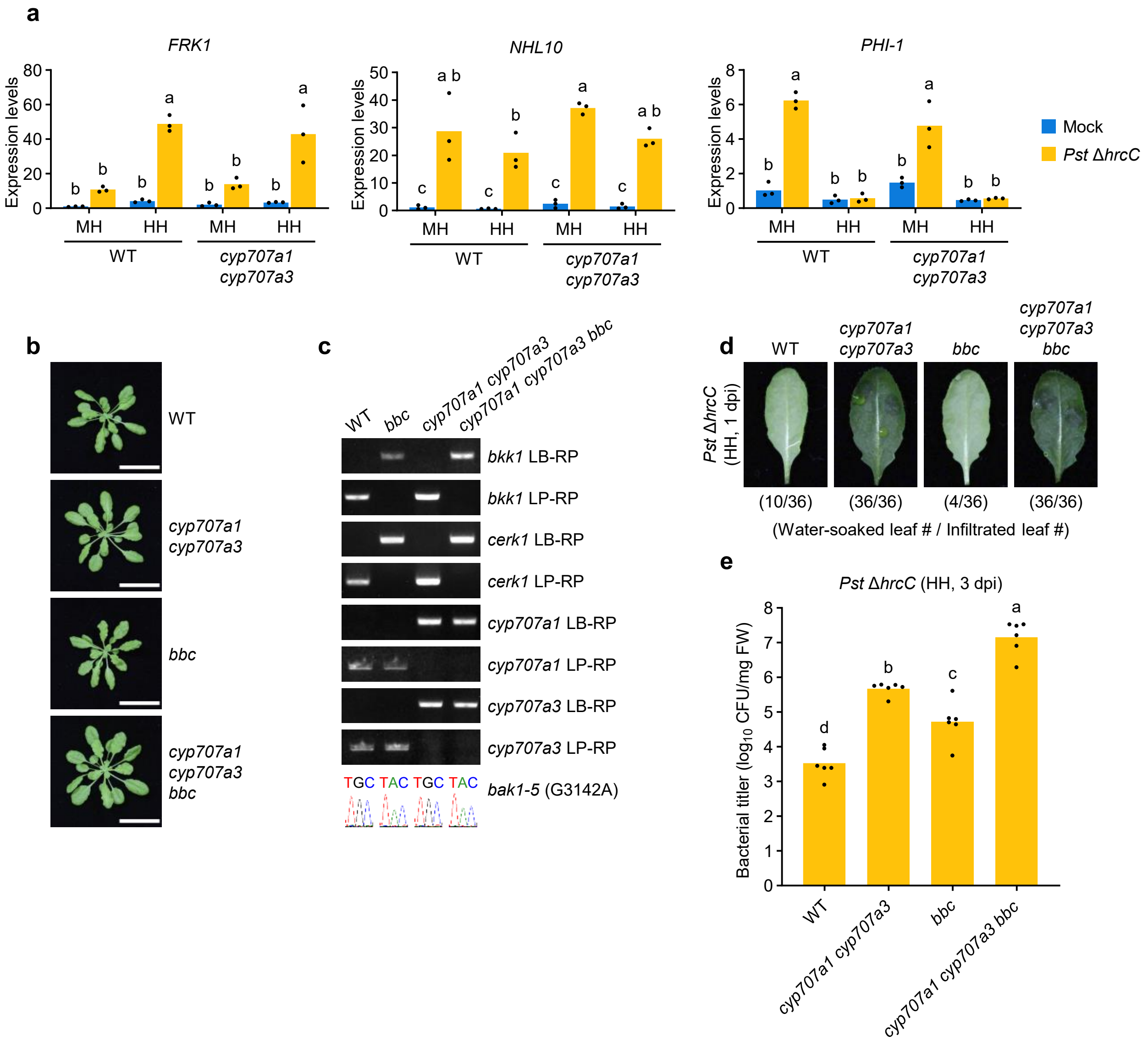


**Supplementary Fig. 4: CYP707A3/A1-mediated ABA depletion and PTI contribute additively to bacterial resistance under high humidity.**

**a**, Expression levels of *FRK1*, *NHL10*, and *PHI-1* in *Pst* Δ*hrcC*-infiltrated leaves under different humidity conditions. The indicated plants were maintained under moderate humidity (MH) or high humidity (HH) for 24 h before infiltration with water (Mock) or *Pst* Δ*hrcC* ($OD_{600}$ = 0.2). Infiltrated plants were kept under the same humidity conditions for 1 h. Bars represent means ($n$ = 3). **b**, Five-week-old plants grown under short-day conditions. Scale bars = 3 cm. **c**, Genotyping of the indicated plants by PCR and Sanger sequencing. LB-RP and LP-RP primer pairs specifically amplify mutated and non-mutated DNA, respectively. **d,** Water-soaking in *Pst* Δ*hrcC*-infiltrated leaves. The indicated plants were infiltrated with *Pst* Δ*hrcC* ($OD_{600}$ = 0.02) and maintained under HH for 1 day. The number of water-soaked leaves was pooled from two independent experiments. **e**, Bacterial growth in *Pst* Δ*hrcC*-infiltrated leaves. The indicated plants were infiltrated with *Pst* Δ*hrcC* ($OD_{600}$ = 0.02) and maintained under HH for 3 days. Bars represent means ($n$ = 6). Different letters indicate statistically significant differences ($P < 0.05$; two-way ANOVA followed by Tukey's test in **a**; one-way ANOVA followed by Tukey's test in **e**). WT, wild-type; *bbc*, *bak1-5 bkk1 cerk1*; dpi, days post-infiltration. Experiments in (**a**) and (**e**) were repeated twice with similar results.

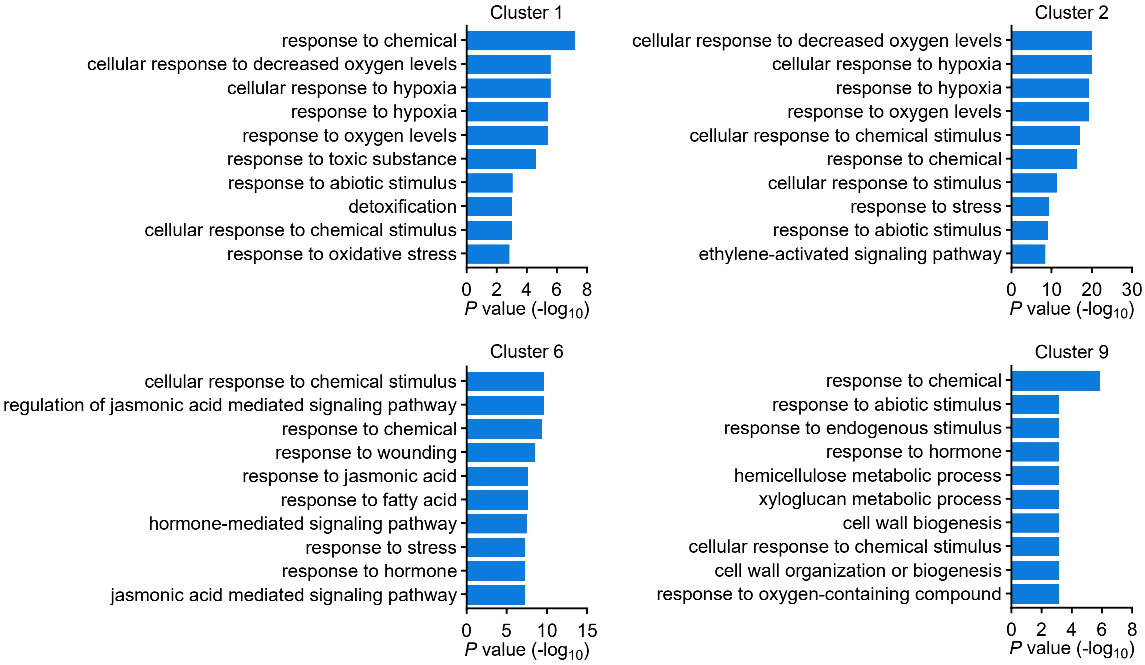


**Supplementary Fig. 5: Gene Ontology analysis of high humidity-induced genes.**
Top 10 enriched Gene Ontology categories of humidity-induced genes in the clusters identified in Fig. 3c.

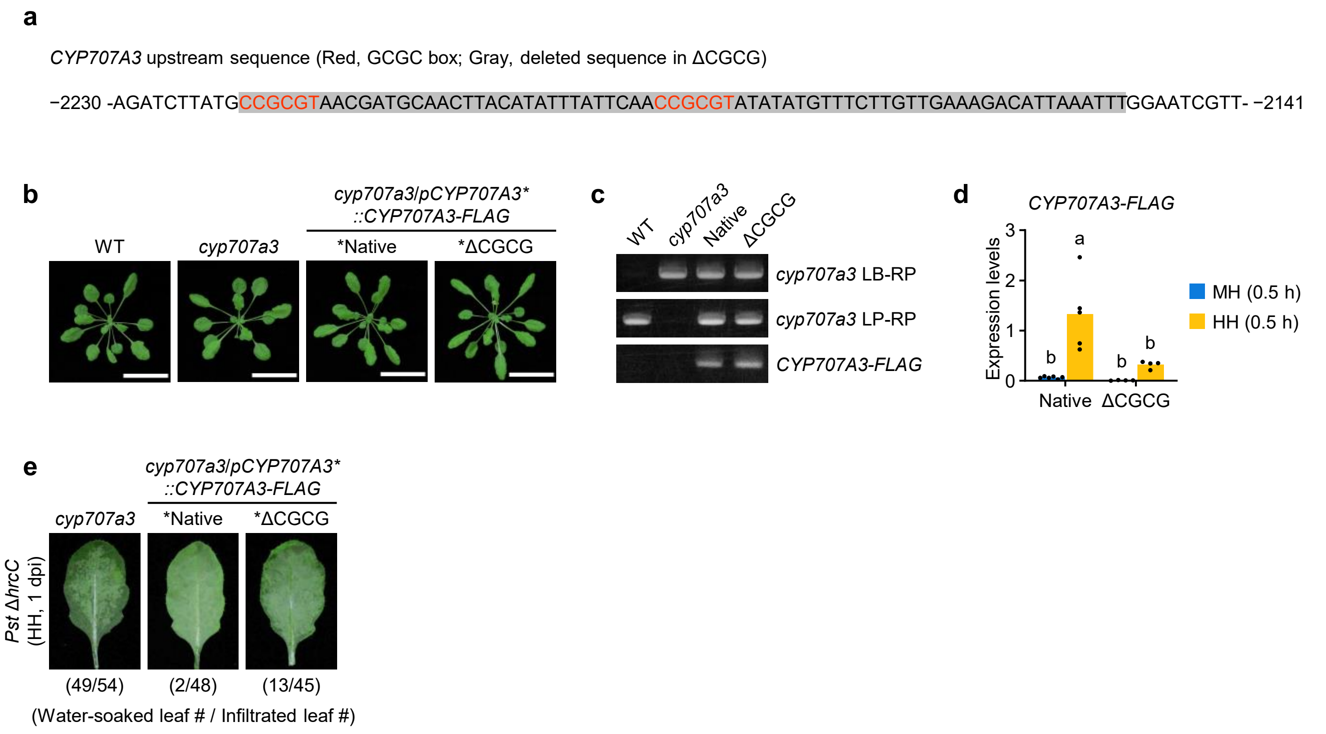


**Supplementary Fig. 6: The CGCG boxes in the *CYP707A3* promoter are critical for water-soaking resistance.**

**a**, Sequence of the *CYP707A*3 promoter region showing the deleted CGCG boxes in the ΔCGCG construct. Numbers indicate the position relative to the ATG start codon. **b**, Five-week-old plants grown under short-day conditions. Scale bars = 3 cm. **c**, Genotyping of the indicated plants by PCR. LB-RP and LP-RP primer pairs specifically amplify mutated and non-mutated DNA, respectively. **d**, Expression levels of *CYP707A3-FLAG* in humidity-treated leaves. The indicated plants were exposure to moderate humidity (MH) or high humidity (HH) for 0.5 h. Bars represent means ($n$ = 3). Different letters indicate statistically significant differences ($P < 0.05$; two-way ANOVA followed by Tukey's test). **e**, Water-soaking in *Pst* Δ*hrcC*-inoculated leaves. The indicated plants were infiltrated with *Pst* Δ*hrcC* ($OD_{600}$ = 0.02) and maintained under HH for 1 day. The number of water-soaked leaves was pooled from three independent experiments. WT, wild-type; dpi, days post-infiltration.

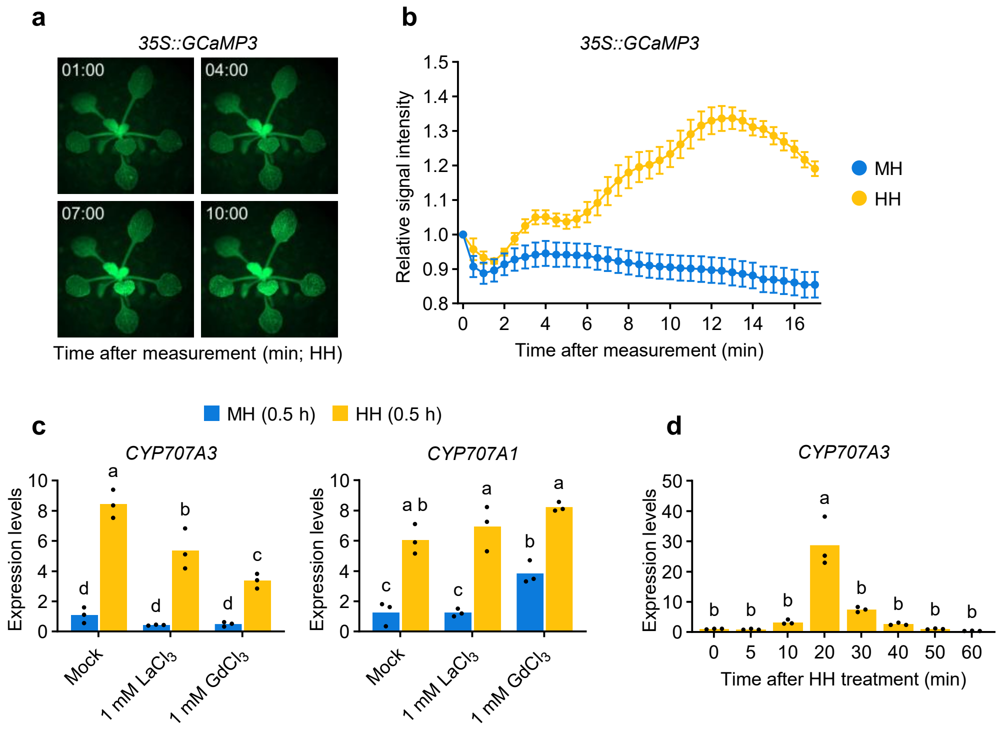


**Supplementary Fig. 7: High humidity-triggered cytosolic $Ca^{2+}$ elevation precedes *CYP707A3* induction.**

**a**, **b,** Cytosolic $Ca^{2+}$ levels in humidity-treated leaves. Two-week-old *35S::GCaMP3* transgenic plants grown under long-day conditions were exposed to moderate humidity (MH) or high humidity (HH) for the indicated time, and GCaMP3 fluorescence signals were measured. Representative GCaMP3 fluorescence images from HH-treated plants are shown in (**a**), and quantified GCaMP3 fluorescence signals are presented in (**b**). Bars represent means ± SEM ($n$ = 10). **c**, Effect of $Ca^{2+}$ channel inhibition on high humidity-induced *CYP707A3* and *CYP707A1* expression. Wild-type (WT) plants were infiltrated with the indicated inhibitors, maintained under MH for 24 h, and then exposed to MH or HH for 0.5 h. Bars represent means ($n$ = 3). **d**, Temporal dynamics of *CYP707A3* expression in high humidity-treated shoots. Two-week-old WT plants grown under long-day conditions were exposed to HH for the indicated time. Bars represent means ($n$ = 3). Different letters indicate statistically significant differences ($P$ < 0.05; two-way ANOVA followed by Tukey's test in **c**; one-way ANOVA followed by Tukey's test in **d**). Experiments in (**c**) and (**d**) were repeated twice with similar results.

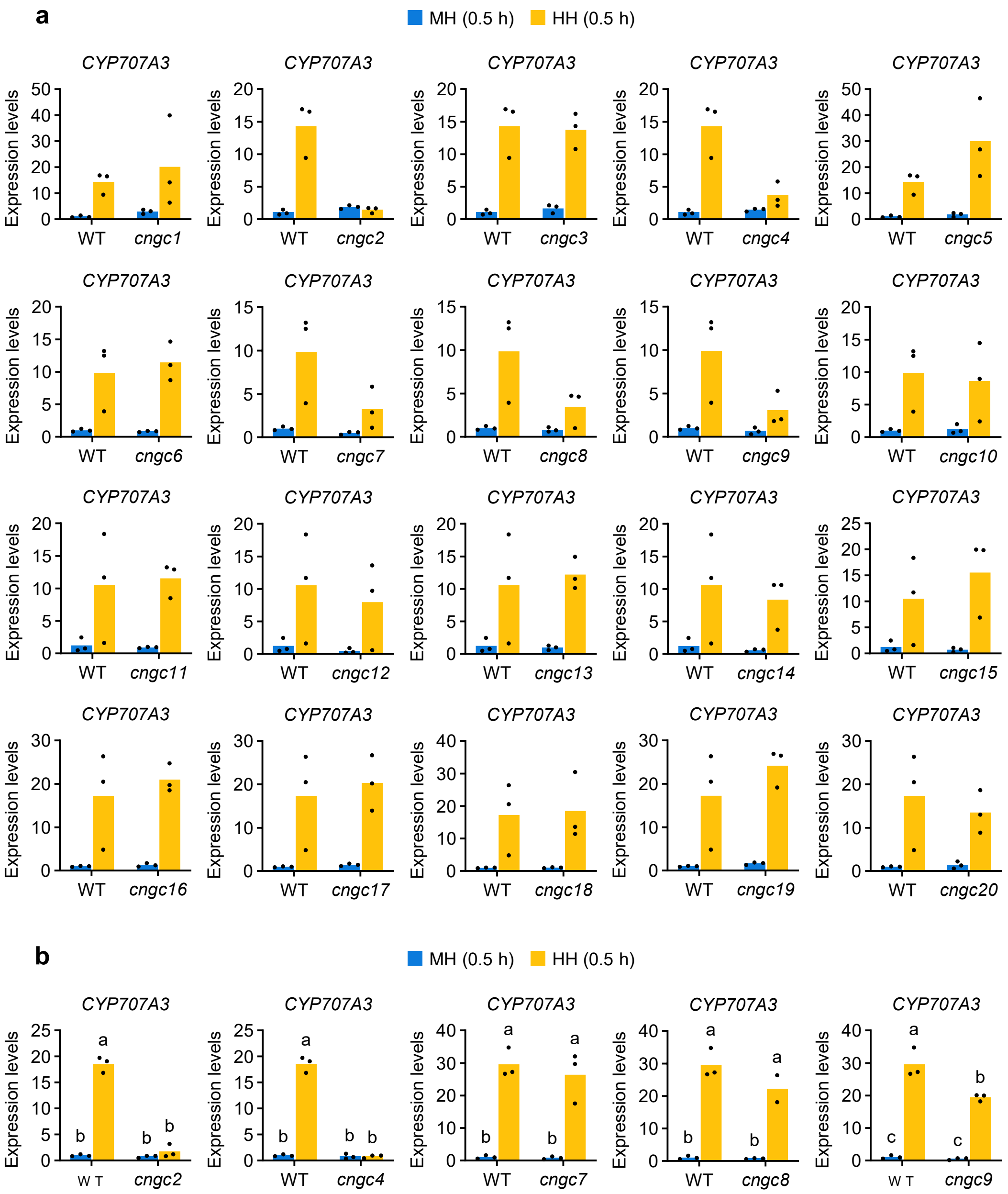


**Supplementary Fig. 8: High humidity-induced *CYP707A3* expression is impaired in *cngc2*, *cngc4*, and *cngc9* mutant plants.**

**a**, **b**, Expression levels of *CYP707A3* in humidity-treated shoots. The indicated plants were grown under long-day conditions for 2 weeks and then exposed to moderate humidity (MH) or high humidity (HH) for 0.5 h. Results from the first experiments are shown in (**a**), and results from the second experiments are shown in (**b**). Bars represent means ($n$ = 2-3). Different letters indicate statistically significant differences ($P < 0.05$; two-way ANOVA followed by Tukey's test). WT, wild-type.

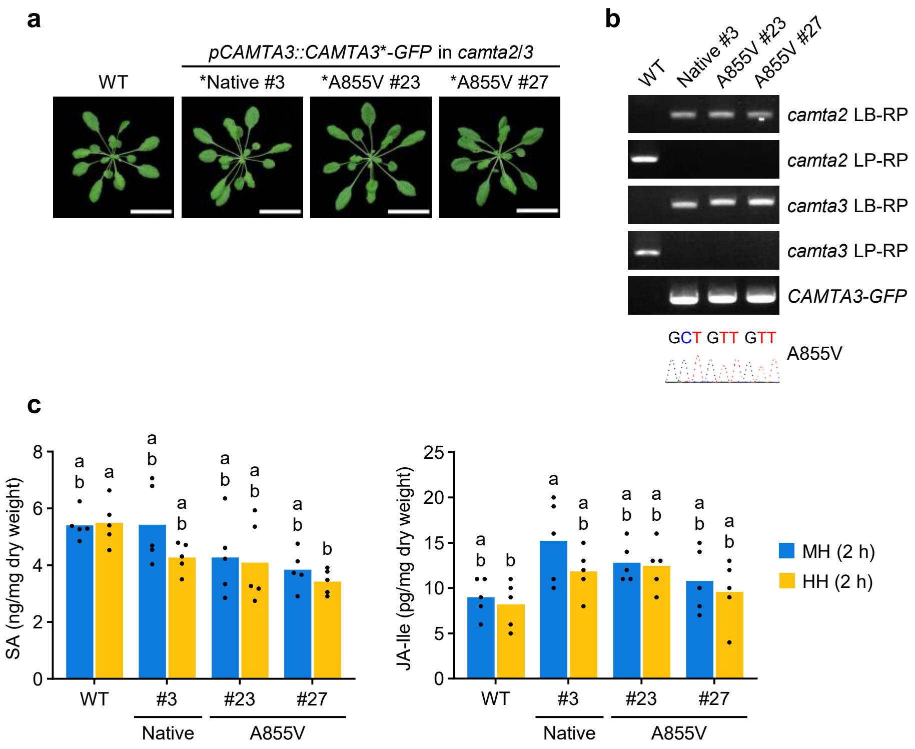


**Supplementary Fig. 9: Characterization of *camta3* mutant plants.**
**a**, Five-week-old plants grown under short-day conditions. Scale bars = 3 cm. **b**, Genotyping of the indicated plants by PCR and Sanger sequencing. LB-RP and LP-RP primer pairs specifically amplify mutated and non-mutated DNA, respectively. **c**, SA and JA-Ile levels in humidity-treated leaves. The indicated plants were exposed to moderate humidity (MH) or high humidity (HH) for 2 h. Bars represent means ($n$ = 5). Different letters indicate statistically significant differences ($P < 0.05$; two-way ANOVA followed by Tukey's test). WT, wild-type; CAMTA3-A855V, a partial loss-of-function variant.

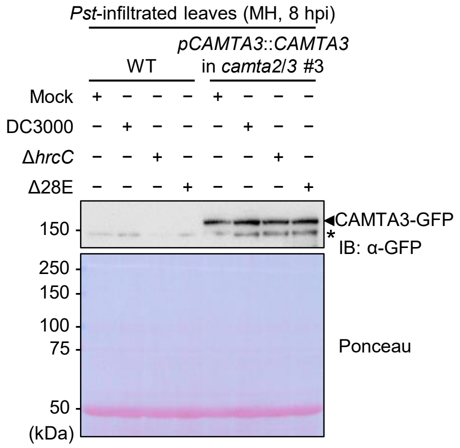


**Supplementary Fig. 10: CAMTA3 protein levels in *Pst* DC3000-inoculated leaves.**
Wild-type (WT) and transgenic plants expressing CAMTA3-GFP driven by the *CAMTA3* promoter (*pCAMTA3::CAMTA3* in *camta2/3* #3) were infiltrated with the indicated *Pst* DC3000 strains ($OD_{600}$ = 0.02) and maintained under moderate humidity (MH) for 8 h. CAMTA3-GFP was detected by immunoblotting with anti-GFP antibody (α-FLAG). The ponceau-stained blot is shown as a loading control. The experiment was repeated three times with similar results.

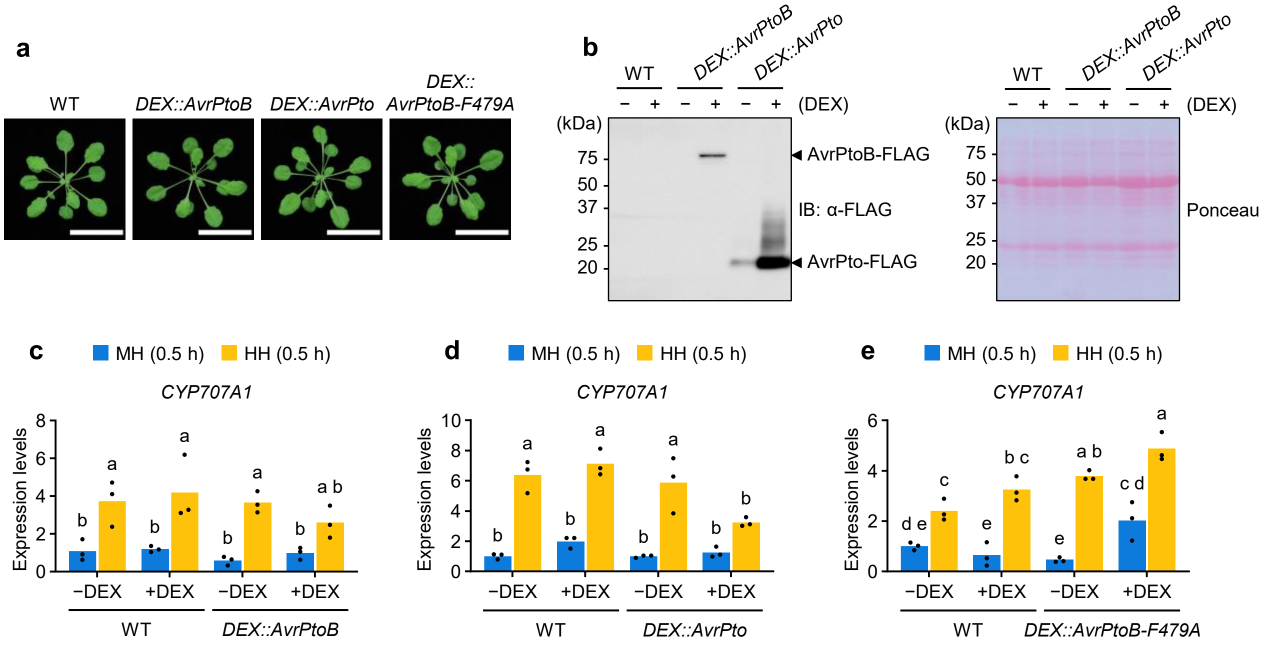


**Supplementary Fig. 11: Characterization of *DEX::AvrPtoB*, *DEX::AvrPto*, *and DEX::AvrPtoB* transgenic plants.**

**a**, Five-week-old plants grown under short-day conditions. Scale bars = 3 cm. **b**, Expression levels of AvrPtoB-FLAG and AvrPto-FLAG in DEX-treated leaves. The indicated plants were infiltrated with 0.1% ethanol (−DEX ) or 10 µM DEX (+DEX) and maintained under moderate humidity (MH) for 24 h. AvrPtoB-FLAG and AvrPto-FLAG were detected by immunoblotting with anti-FLAG antibody (α-FLAG). The ponceau-stained blot is shown as a loading control. **c-e**, Expression levels of *CYP707A1* in humidity-treated leaves with or without *AvrPtoB* (**c**), *AvrPto* (**d**), and *AvrPtoB-F479A* (**e**). The indicated plants were treated with DEX as described in (**b**), followed by exposure to MH or high humidity (HH) for 0.5 h. Bars represent means ($n$ = 3). Different letters indicate statistically significant differences ($P$ < 0.05; two-way ANOVA followed by Tukey's test). The experiments in (**c**) and (**d**) were repeated twice with similar results.

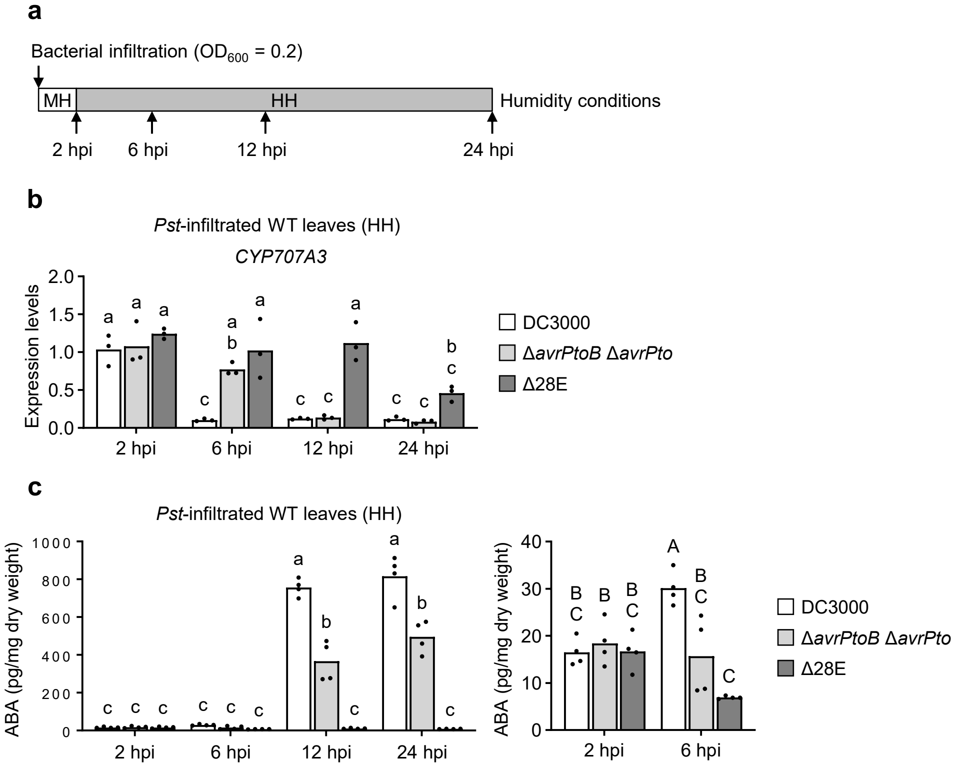


**Supplementary Fig. 12: *Pst* DC3000-mediated *CYP707A3* suppression and ABA accumulation are compromised by loss of *avrPtoB* and *avrPto***

**a**, Schematic overview of the experimental design for monitoring temporal dynamics of *CYP707A3* expression and ABA accumulation in *Pst* DC3000-inoculated leaves under high humidity (HH). **b**, Temporal dynamics of *CYP707A3* expression in *Pst* DC3000-inoculated leaves. Wild-type (WT) plants were infiltrated with the indicated *Pst* DC3000 strains ($OD_{600}$ = 0.2), maintained under moderate humidity (MH) for 2 h (2 hours post-infiltration, hpi), and then transferred to HH for 4 h (6 hpi), 10 h (12 hpi), or 22 h (24 hpi). Bars represent means ($n$ = 3). **c**, Temporal dynamics of ABA levels in *Pst* DC3000-inoculated leaves. WT plants were infiltrated with the indicated *Pst* DC3000 strains as described in (**a**) and (**b**). Bars represent means ($n$ = 4). Different letters indicate statistically significant differences ($P$ < 0.05; two-way ANOVA followed by Tukey's test). Experiment in (**b**) was repeated twice with similar results.

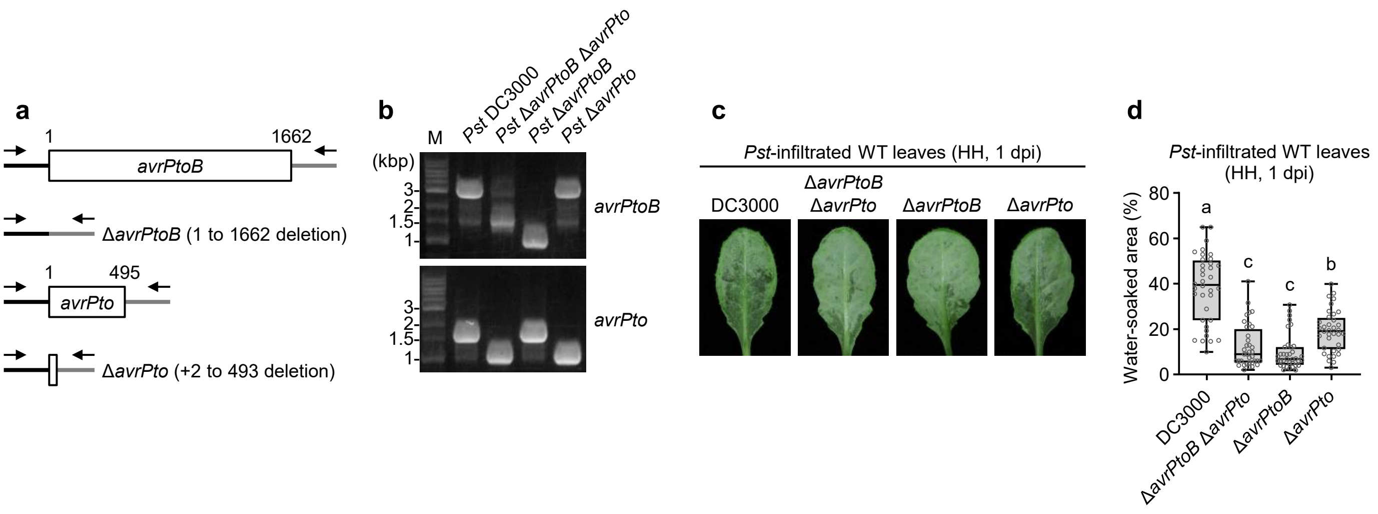


**Supplementary Fig. 13: Characterization of *Pst* Δ*avrPtoB* and *Pst* Δ*avrPto* mutant strains.**
**a, b**, Generation of *Pst* DC3000 mutant strains lacking AvrPtoB (Δ*avrPtoB*) or AvrPto (Δ*avrPto*). Deleted genomic regions (**a**) and genotyping results (**b**) of the established *Pst* DC3000 mutants. Numbers indicate the position relative to the ATG start codon. Arrows indicate genotyping primers. M, DNA size marker. **c**, **d**, Representative images (**c**) and quantification (**d**) of water-soaking in *Pst* DC3000-inoculated leaves. Wild-type (WT) plants were infiltrated with the indicated *Pst* DC3000 strains ($OD_{600}$ = 0.2), maintained under MH for 2 h, and then transferred to HH for 20-22 h. Bars represent the median ($n$ = 36; pooled from two independent experiments). Different letters indicate statistically significant differences ($P$ < 0.05; one-way ANOVA followed by Tukey's test).
